# Visualization of mitochondrial cristae and mtDNA evolvement and interactions with super-resolution microscopy

**DOI:** 10.1101/2022.12.26.521907

**Authors:** Wei Ren, Xichuan Ge, Meiqi Li, Shiyi Li, Chunyan Shan, Baoxiang Gao, Peng Xi

## Abstract

Mitochondrial cristae host the respiratory chain complexes composed of mitochondrial DNA (mtDNA)-encoded and nuclear-encoded proteins and are responsible for ATP production. Movement of mtDNA located in the matrix is limited due to blockade by the cristae; yet, the dynamic interaction between the inner membrane and mtDNA remains unknown due to the insufficient spatiotemporal resolution of conventional microscopy and the lack of appropriate *in vivo* probes targeted to the mitochondrial inner membrane. Here, we developed a novel fluorescence probe to visualize the inner membrane using low-power stimulated emission depletion (STED) microscopy. Dual-color imaging of the inner membrane and mtDNA demonstrated that mtDNA is more likely to spread at mitochondrial tips or branch points under an overall even distribution. Interestingly, exploration of forming this distribution propensity uncovered that the mitochondrial dynamics are closely related to the location of mtDNA, and further insight found that fusion always occurs near mtDNA in order to minimize the pressure for cristae remodeling. In healthy cells, mitochondrial dynamics based on cristae remodeling promotes the even distribution of mtDNA, on the contrary, when cristae structure fails in apoptosis and ferroptosis, leading to mtDNA distribution disorder. Observation of active changes during apoptosis further captured the dynamic process of inner membrane herniation and mtDNA leakage along with cristae remodeling. Under ferroptosis, the mitochondria shrank into ellipsoids and mtDNA converged at the center of mitochondria. The rich dynamics between the cristae and mtDNA, revealed at unprecedented spatiotemporal resolution, show the motive and outgrowth of mtDNA distribution.

## Introduction

Mitochondria are highly dynamic organelles that orchestrate a vast range of biological processes from the production of ATP to the biosynthesis of iron-sulfur clusters, calcium homeostasis, and various cellular signaling [1–5]. Mitochondria, as double-membrane organelles, are structurally divided into four regions: outer membrane (OM), inner membrane (IM), intermembrane space, and matrix. The IM can be further divided by the cristae junction (CJ) into the inner boundary membrane, juxtaposed with OM, and the cristae, which are deeply convoluted invaginations that provide expansion of the surface area [6, 7]. In response to different physiological or pathological conditions, mitochondria regulate cristae biogenesis or reorganize cristae structures with cooperation from mitochondrial-shaping proteins [8]. Perturbations of mitochondrial-shaping proteins derange cristae structure, sway the OXPHOS system, and damage cellular metabolism [6, 9]. Mitochondrial DNA (mtDNA) contains 37 genes and encodes 13 mitochondrial proteins that are incorporated into mitochondrial respiratory chain complexes and utilized in oxidative phosphorylation (OXPHOS) system for the production of ATP [10], so the maintenance of mtDNA distribution is essential for mitochondrial function. MtDNA is packaged by proteins to form nucleoids, which are approximately 100 nm in size and are distributed in the void matrixes between cristae clusters to perform functions [11–13].

Once replication is complete, mtDNA must be spread throughout the mitochondrial network. Because the OXPHOS complexes consist of mtDNA-encoded and nuclear-encoded subunits, they are assembled *in situ,* neighboring to the nucleoid [10, 14]. Movement of mitochondrial nucleoids located in the matrix compartment is limited due to the fact that mitochondrial cristae represent a substantial barrier to longitudinal free diffusion [13]; there are also opinions for active trafficking of nucleoids under certain circumstances[15, 16]. Regardless, mitochondrial fusion and fission involved in cristae remodeling are essential for the distribution and maintenance of mtDNA [17, 18], and dynamic cristae are also required to facilitate the normal rate of fission and fusion [10]. Recent evidence shows that deficiencies in the mitochondrial contact site and cristae organizing system (MICOS) complex result in cristae disconnection, abnormal distribution of nucleoids, and OXPHOS defects, suggesting that the mitochondrial IM plays an important role in the organization, segregation, and distribution of nucleoids [19]. However, the manner by which the dynamic properties of the IM and cristae arrangement facilitate the distribution of mtDNA across the network has been relatively underexplored due to the diffraction limit in optical microscopy and lack of appropriate *in vivo* markers of the mitochondrial IM. Mitochondria form a huge network by fusing or emerging the new branch [20], and the junctions of different branches are the core of the whole network. It is also unknown how the cristae and mtDNA are arranged at these branch point.

The development of super-resolution imaging techniques has made it possible to visualize mitochondrial cristae [21–27]. Single-molecule localization (SML)-based techniques such as PALM/STORM offer high spatial resolution, but the poor temporal resolution cannot effectively observe cristae dynamics [24]. Structured illumination microscopy (SIM) can obtain a resolution of approximately 90–120 nm through image post-processing but is insufficient for observing the dynamics of a single crista and also requires extensive expertise to identify artifacts [25]. By contrast, the spatial and temporal resolution of stimulated emission depletion (STED) microscopy is approximately 50 nm and 1 s, making it the most promising tool for dynamic imaging of mitochondrial cristae. Nevertheless, utilization of STED is limited in mitochondria because probe photobleaching occurs during time-lapse live-cell imaging. Various measures have been introduced to minimize photobleaching in STED nanoscopy, including the development of photostable fluorophores [28], the use of fluorophores with multiple off-states [29], and the exchange of fluorophore labels with a wash-free imaging buffer reservoir [30]. In addition, a prerequisite for the super-resolution imaging of mitochondrial cristae is the targeted accumulation of dyes in the mitochondrial IM [31]. Moreover, since STED requires a high intensity depletion beam for resolution enhancement but mitochondria are very sensitive to light stimulation, it poses a new challenge for dyes, which should have both high brightness and a high stimulated emission cross-section. Here, we developed a fluorescent dye, IMMBright660, which has the characteristics of outstanding photostability, selective labeling of the mitochondrial IM, and emission of light only upon binding to the IM. Using STED nanoscopy with IMMBright660, we were able to record mitochondrial cristae dynamics with 40-nm spatial resolution and an observation window of more than 500 frames.

## Results

### A novel STED probe targeting the inner mitochondrial membrane

We developed a mitochondrial IM-localized probe named IMMBright660, with the chemical structure shown in Fig. 1a. Si-rhodamine compounds are far-red mitochondrial dyes with high fluorescence quantum yields, good photostability, and excellent membrane permeability; however, these cationic dyes only accumulate in the mitochondrial matrix. Considering the fact that the tight bilayer of the mitochondrial IM is mainly composed of phospholipids, we hypothesized that highly selective mitochondrial IM dyes can be designed by introducing a lipophilic alkyl chain to Si-rhodamine dyes; thus, we prepared IMMBright660 by modifying Si-rhodamine with n-octanoic acid (Fig. 1a). IMMBright660 exhibited high photostability, chloride ion-induced fluorescence quenching, lipid membrane-activated fluorescence emission characteristics, and mitochondrial IM localization, allowing us to perform wash-free STED imaging (Fig. S1-S3). The excitation and emission peaks of IMMBright660 are at 660 nm and 680 nm, respectively (Fig. 1b). The spectroscopic properties of IMMBright660 are compatible with STED using NIR 775-nm depletion lasers. In commercial STED, near-infrared depletion lasers are advantageous since they cause less damage to biological objects and benefit from reduced autofluorescence, photobleaching, and light scattering. We measured the saturation intensity *I*_sat_ of IMMBright660, which was directly related to the resolution of STED imaging. IMMBright660 showed a very low saturation intensity of 0.864 mW (Fig. S4), suggesting that IMMBright660 is an excellent STED dye for the mitochondrial IM. The stimulated emission cross-section is related to the emission spectrum of the dye [32, 33]. The excitation and emission spectrum of IMMBright660 have an overall redshift, leading to lower saturation power. The effect of secondary excitation on imaging can be ignored (Fig. S5) [33].

**Figure 1.**
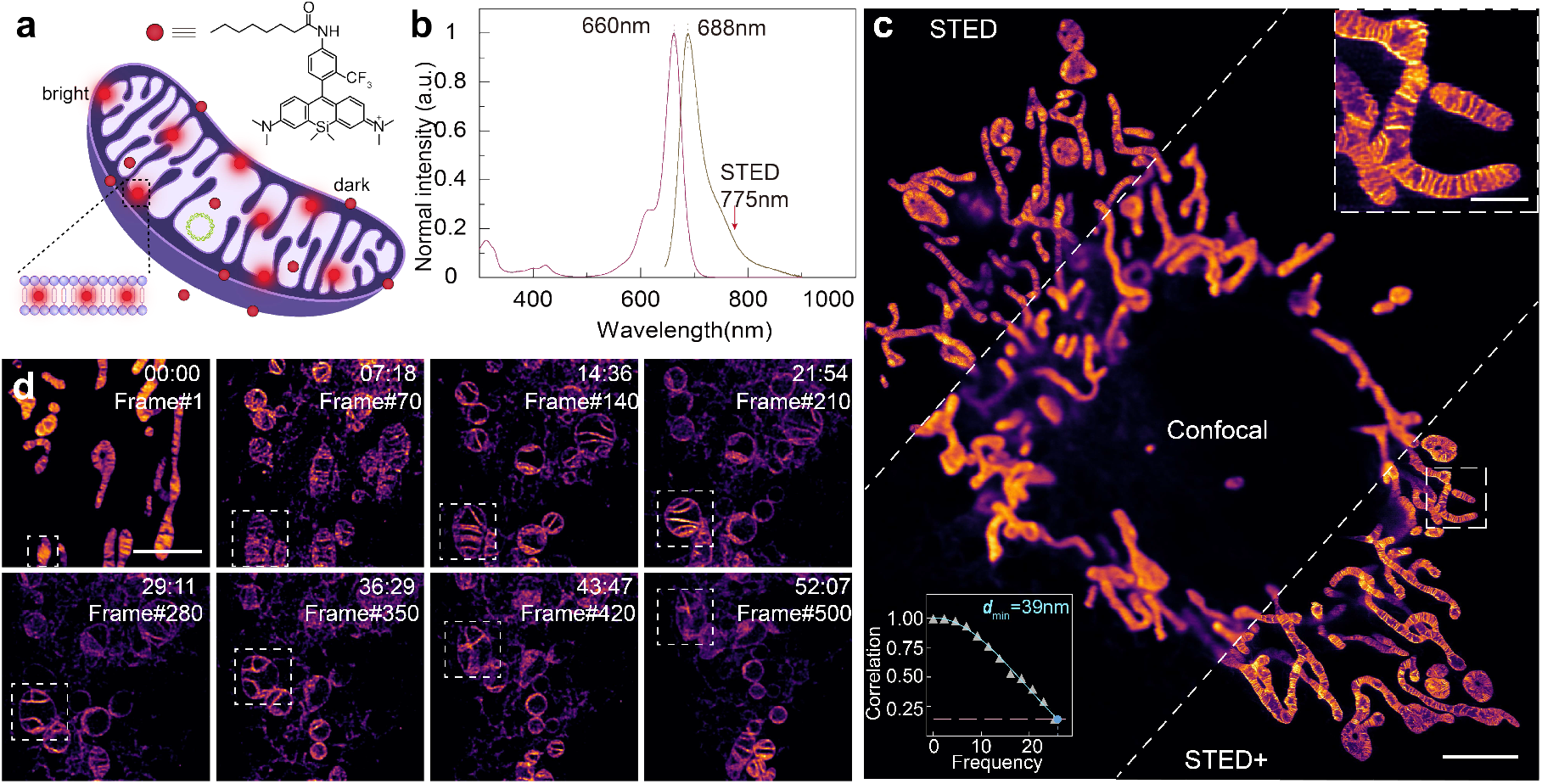
A novel probe IMMBright660 for long-time mitochondrial STED imaging. **a**, Chemical structure of IMMBright660 used for the specific labeling of the mitochondrial inner membrane. **b**, The absorption and emission spectra of IMMBright660, which can be depleted using a 775-nm laser. **c**, Comparison of confocal, STED, and STED+ (deconvolution with Huygens) imaging results in living COS7 cell mitochondria labeled with IMMBright660. The lower left corner shows the Fourier ring correlation (FRC) calculation for STED+. The upper right corner shows the enlarged results for STED+. **d**, A long period time-lapse STED imaging of mitochondria. White dotted boxes indicate the same mitochondrion. Scale bars: in **c**, 5 μm; enlarged image in **c**, 1 μm; and in **d**, 3 μm.

As a fluorogenic probe, IMMBright660 allows us to perform wash-free STED experiments, since it is insensitive to photobleaching and suitable for long-term STED. Firstly, we implemented single-frame wash-free live-cell imaging using IMMBright660-labeled COS7 cells, and the results show that mitochondrial cristae could be clearly resolved under STED and STED+ in comparison with confocal (Fig. 1c). The absolute resolution of STED and STED+ obtained by Fourier ring correlation (FRC) reached 40 nm and 39 nm [34], respectively, and the full width at half maximum was fitted to 44 nm and 39 nm (Origin) (Fig. S6), respectively.

Subsequently, we used IMMBright660 to perform low-power time-lapse STED imaging to visualize the dynamic changes in mitochondrial internal spatial structure. Under wash-free conditions, we could clearly identify single cristae and monitor their dynamics for more than 500 frames, for a total period of 52 min (Fig. 1d and Movie S1). The excellent stability of IMMBright660 and the reliability of the live-cell STED imaging strategy established a reliable method to study the manner by which mitochondrial cristae membrane dynamics affect the distribution of mtDNA.

### Propensity of mtDNA distribution in the mitochondrial network

All proteins encoded by mtDNA are subunits of OXPHOS enzymes located in the cristae membrane [10]; thus, proper mtDNA distribution within mitochondrial networks are essential for maintaining mitochondrial functions (supply of ATP). We imaged the IM under STED with IMMBright660 labeling and mtDNA under confocal with SYBR™ Gold nucleic acid gel dye (Thermo fisher, US, No. S11494) labeling (Fig. 2a). Dual-color imaging (IM using STED and mtDNA using confocal. Unless otherwise specified, all dual-color imaging methods are the same) revealed that mitochondrial cristae are often arranged in clusters, and the nucleoids generally occupy voids in the matrix between cristae clusters (Fig. 2b). In comparison with references [13, 21], we were able to observe mitochondrial cristae at a resolution of 40 nm in living cells. We also found that mtDNA is regularly spaced within mitochondrial networks, preferring to be distributed at the tips and branch points (Fig. 2c).

**Figure 2.**
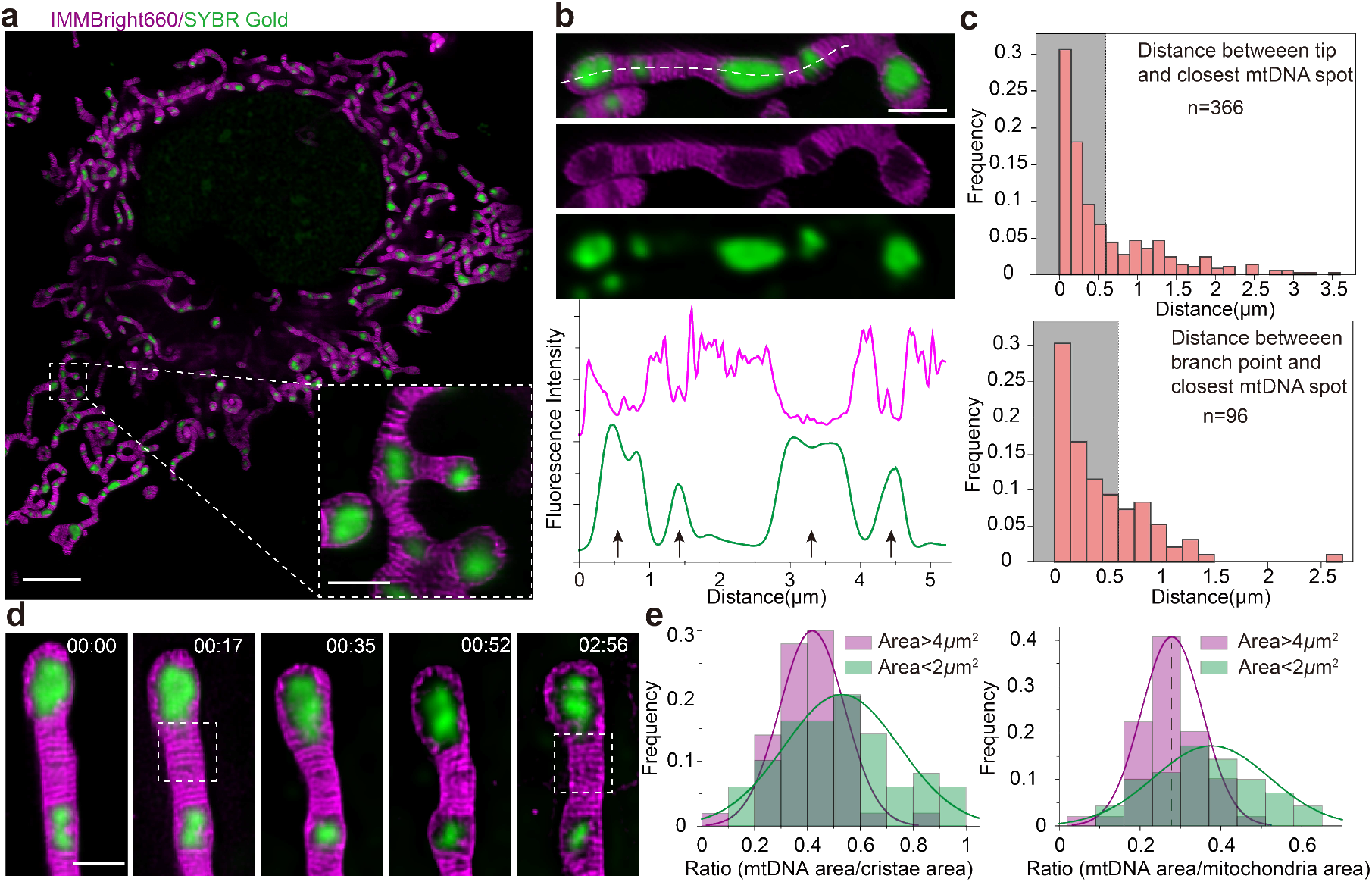
Distribution of cristae and mtDNA in mitochondria. **a**, Dual-color live-cell imaging for mitochondria (STED) and mtDNA (confocal). **b**, The pixel intensity of IMMBright660 and SYBR™ Gold from a line scan drawn along the mitochondria (dashed line); arrows indicate nucleoid positions. **c**, Distance frequency distribution histogram from mitochondrial tips or branch points to the nearest mtDNA. The proportion of the gray part (distance < 0.6 μm) is 65.0% and 67.7%, respectively. **d**, Spatial changes in mtDNA and cristae at the mitochondrial tip by STED imaging. The white boxed area shows cristae remodeling. **e**, Frequency distribution histogram of the ratio of mtDNA area to cristae area or mitochondria area. Scale bars: in **a**, 5 μm; enlarged image in **a**, 1 μm; and in **d**, 0.5 μm.

We then quantitated the propensity of mtDNA distribution and created a frequency distribution histogram by measuring the distances between mitochondrial tips (n = 366) or mitochondrial branch points (n = 96) and the nearest mtDNA (Fig. 2c). Interestingly, these nucleoids near mitochondrial tips or branch points show an exponential distribution, suggesting that nucleoids tend to locate at mitochondrial tips or at branch points of mitochondrial networks [35] (Fig. 2c). Recent evidence shows that after mtDNA replication, (endoplasmic reticulum) ER-linked mitochondrial fission occurs between the replicated mtDNA [36], which also indicates that mtDNA is located at newly generated mitochondrial tips after fission. We dynamically observed the spatial changes in mtDNA and mitochondrial cristae at the tips and found that mitochondrial cristae tended to tether mtDNA to the tips, preventing its movement (Fig. 2d and Movie S2). Thus, the movement of mtDNA is restricted by the cristae at mitochondrial tips.

We examined the difference in mtDNA distribution between small mitochondria and mitochondrial networks by measuring the ratio of mtDNA area to mitochondrial cristae area. In this process, we found that the area ratio (mtDNA area / cristae area) of small mitochondria (0.524) is higher than that of mitochondrial networks or large mitochondria (0.421) (Fig. 2e). We hypothesize that large mitochondria with a low area ratio and dense cristae are used to produce ATP to maintain cellular function, and small mitochondria with a high area ratio can serve as the carriers of mtDNA to achieve even distribution of mtDNA through floating and fusion processes. Cristae are the principal sites of ATP production; therefore, large mitochondria have a lower area ratio than small mitochondria. We further investigated whether this relationship exists over the entire mitochondrial area and found that the area ratio (area mtDNA/area mitochondria) of small mitochondria (0.377) is still higher than that of mitochondrial networks or large mitochondria (0.277) (Fig. 2e). This further illustrates that the area ratio is different in mitochondria of different sizes, and STED also facilitates the area ratio measurement with high accuracy.

### Mitochondrial branch point dynamics associated with mtDNA

The formation and maintenance of the mitochondrial network is critical to the performance of mitochondrial functions (such as respiratory capacity, material exchange, and mtDNA integrity, etc.) [20, 37]. The branch point being at the core of the mitochondrial network, cristae dynamics and mtDNA distribution during the formation of the branch point are yet to be explored, as is the manner by which the distribution of mtDNA affects the dynamic behavior of the branch point. We observed the cristae arrangement and mtDNA distribution at the mitochondrial branch point using dual-color imaging and found that mitochondrial cristae were always arranged perpendicular to the extension direction of the branch regardless of the presence or absence of mtDNA at the branch point, which resulted in a triangular-like arrangement of cristae near the branch point (Fig. 3a, b). From the results in Fig. 2c, it can be found that the probability of the presence of mtDNA at the branch point is 0.677. In the presence of mtDNA, the void matrixes left by the cristae arrangement were filled; however, in the absence of mtDNA, a space was left at the junction.

**Figure 3.**
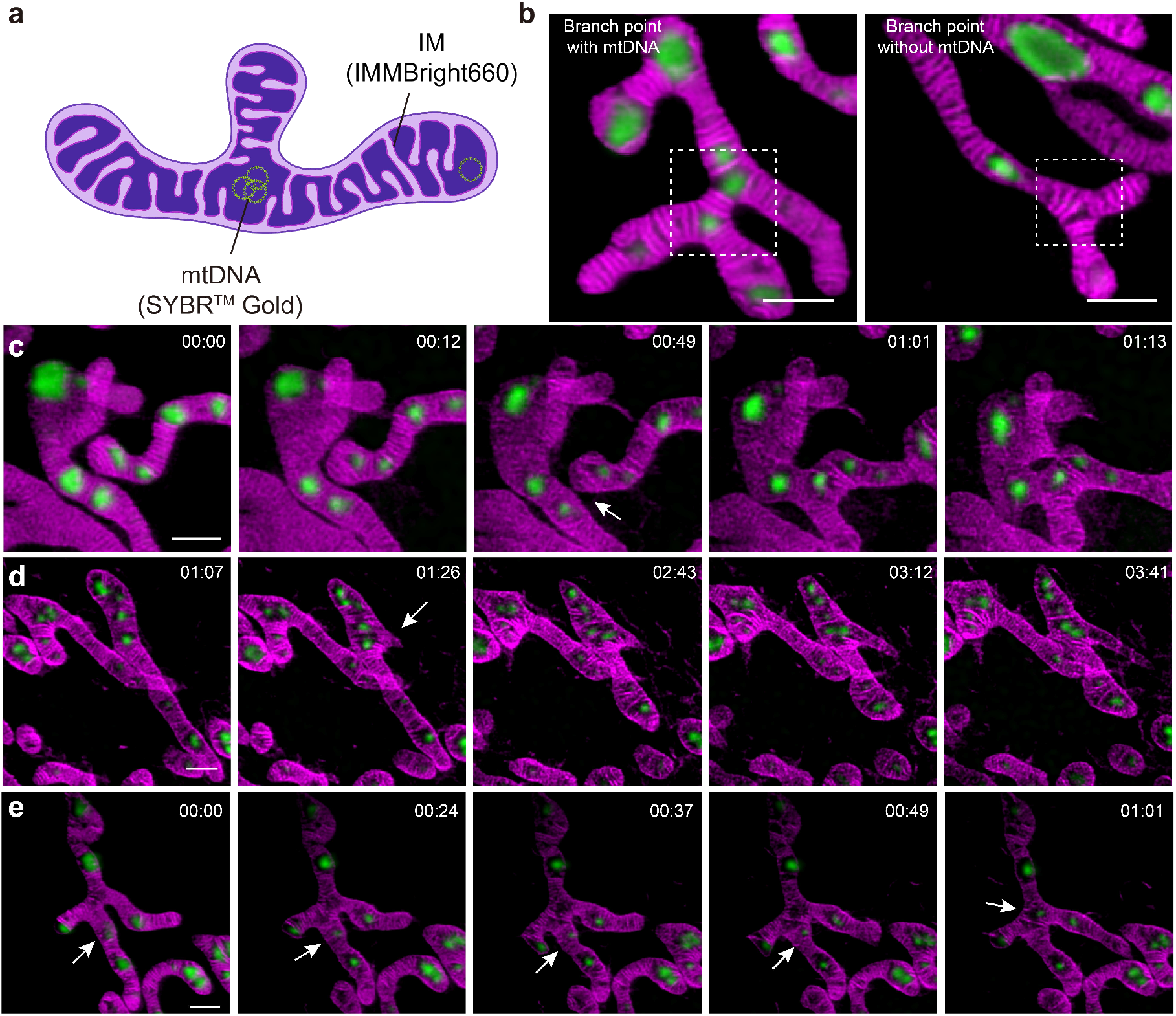
Mitochondrial branch point dynamics associated with mtDNA. **a**, Cartoon showing that mtDNA prefers to be distributed at the tips and branch points. **b**, Presence (left) and absence (right) of mtDNA at branch points; the white box represents the branch point. **c**, Branch point formation by mitochondrial fusion; white arrows indicate fusion sites. **d**, Branch point formation by emersion of a new branch; white arrows indicate from where the new branch was generated. **e**, The mtDNA on the mitochondrial branch moves to the branch point; white arrows indicate the movement of mtDNA. Scale bars in **b**–**e**, 1 μm.

The first way that branch point formation can be achieved is by mitochondrial fusion, with the fusion site becoming a new branch point. As shown in Fig. 3c (Movie S3), the two mitochondria gradually approached one another, and the respective mtDNAs were relatively close to the fusion site. A new branch point was formed after fusion; and since fusion occurred near mtDNA, the mtDNA at the fusion site was naturally at the branch point. The second way that branch point formation can be achieved is by emerging the new branch [16, 20]. We visualized cristae dynamics and mtDNA changes in distribution during this process under STED for the first time, and Fig. 3d (Movie S4) shows the entire process of mitochondrial tubule extension to form branches. Mitochondria extended a new branch from a spatially loose mtDNA site. As the length and width of the extended branch increased, the mitochondrial cristae at the extension site were rapidly remodeled to adapt to the new branch. Since the number of cristae at the newly generated branch point were reduced, the mtDNA activity range increased, and moreover, the cristae at the newly generated mitochondrial branch were relatively sparse.

Since the formation of both branch points occurs near mtDNA, mtDNA is located near the newly formed branch point. Due to the regularity of cristae arrangement at the branch point, void matrixes will be left when there is no mtDNA at the branch point. In order to efficiently utilize the limited space, mtDNA will actively move to the junction. As shown in Fig. 3e (Movie S5), mtDNA was initially located at the mitochondrial branch; but over time, as indicated by the white arrow, moved to the branch point. In the presence of mtDNA, void matrixes were formed between the cristae clusters; however, in the absence of mtDNA, the void matrixes between the cristae clusters gradually disappeared.

### Mitochondrial membrane dynamics promote the even distribution of mtDNA during fusion and fission

In comparison with other organelles (e.g., peroxisomes and lysosomes), mitochondria cannot be created *de novo,* and their mtDNA must be replicated in order to be transmitted to daughter mitochondria. The movement of nucleoids within mitochondria is restricted; thus, nucleoid distribution rely on mitochondrial fusion and fission processes [17]. It has been observed that ER–mitochondria contact sites are typically located near the replicated nucleoids, indicating that fission is intrinsically related to mtDNA segregation [36]. Fusion is required for proper inheritance and maintenance of mtDNA, maintenance of mtDNA replication and distribution, and dilution of mutated mtDNA [37, 38]. Mitochondrial fission and fusion are critical for achieving even distribution of mtDNA. To study the manner by which mitochondrial membrane dynamics promote the even distribution of mtDNA, we first observed the dynamic changes in cristae and mtDNA during fusion (Fig. 4a and Movie S6) and found that fusion occurred near a pair of mtDNAs between parallel mitochondria (white box). During the fusion process, the two mitochondria moved closer to one another and the IM was remodeled, which was followed by the transmission of mtDNA. A cartoon corresponding to this process is shown in Fig. 4b. Thus, mitochondrial cristae facilitate the transmission of mtDNA between different mitochondria through a series of remodeling processes.

**Figure 4.**
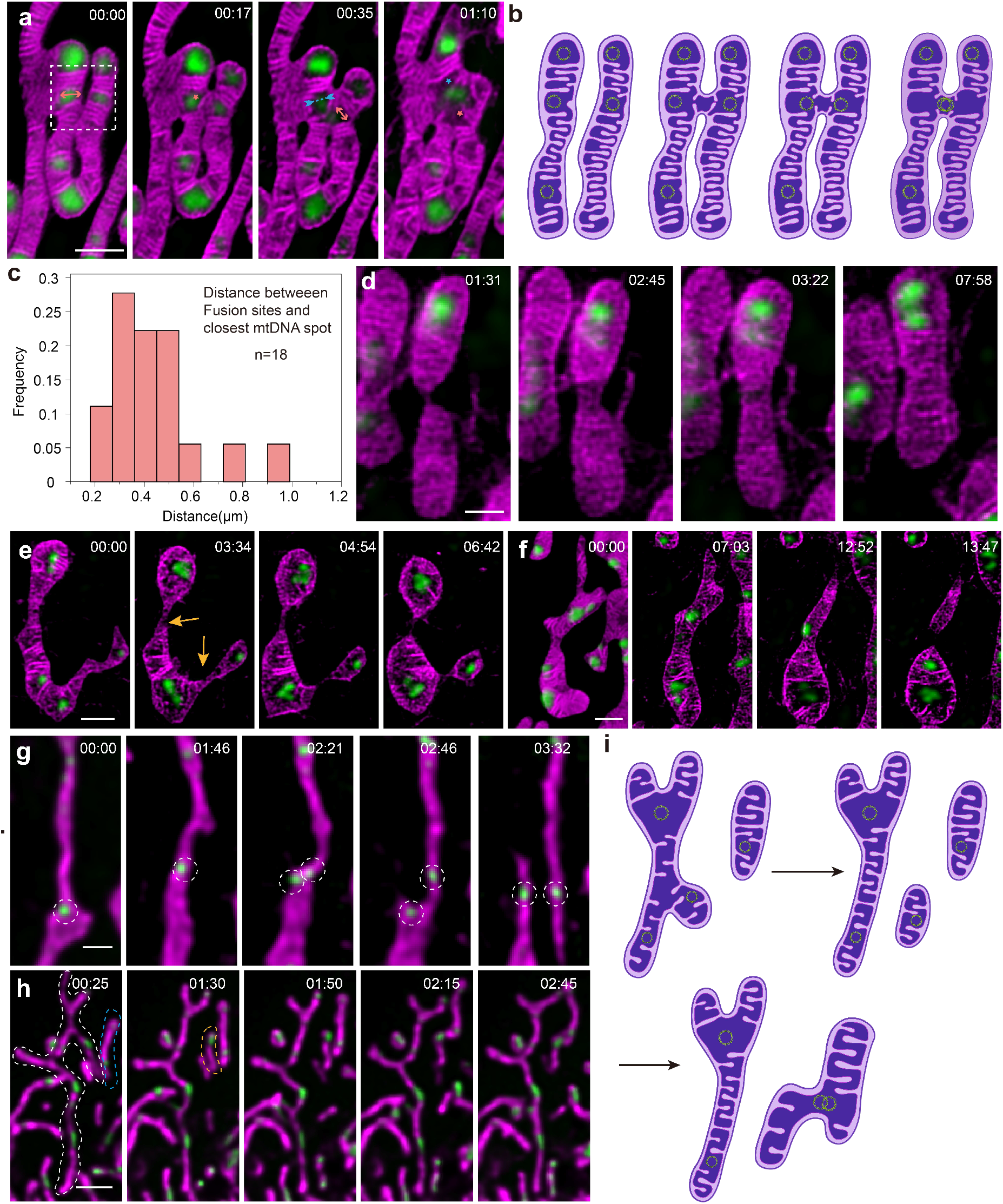
mtDNA distribution and cristae dynamics during mitochondrial fusion and fission. **a**, Fusion process of mitochondria. Merging and splitting events of cristae are indicated by blue and orange asterisks, respectively, with bidirectional arrows facing inward (blue) and outward (orange) representing impending cristae merging and splitting events, respectively. **b**, Cartoon showing the fusion process corresponding to the white box in Figure 1a. **c**, Distance frequency distribution histogram. **d**, Fusion of one mitochondrion containing mtDNA and another mitochondrion without mtDNA. **e**, Fission process of mitochondria. The yellow arrow indicates the fission site. **f**, Mitochondria after fission may not contain mtDNA. **g**, mtDNA replication initiates mitochondrial division. The dotted line shows the replication of mtDNA. **h**, Small mitochondrial branches detach from the mitochondrial network and fuse with nearby mitochondria. White and blue dashed lines represent the mitochondrial network before fission and mitochondria before fusion, respectively. The yellow dotted line represents the small mitochondria after division. **i**, Cartoon corresponding to **h**. Scale bars: in **a** and **e**–**g**, 1 μm; in **d**, 0.5 μm; and in **h**, 2 μm.

We observed the dynamic process of mitochondrial fusion in several experiments and found that fusion always occurs near mtDNA. We subsequently measured the Euclidean distance between the fusion site and the nearest mtDNA (n = 18) to obtain a distance frequency distribution histogram showing an exponential distribution (Fig. 4c), which suggests that the fusion site is extremely close to the mtDNA. The void matrixes between cristae clusters where mtDNA is located are not dense and the cristae arrangement is loose; thus, we hypothesized that the pressure of cristae remodeling will be less when fusion occurs near mtDNA.

We observed that one mitochondrion with two nucleoids fused with another mitochondrion without nucleoids. Interestingly, in comparison with mitochondria without mtDNA, newly fused mitochondria have more abundant cristae through cristae remodeling, suggesting that mitochondrial function can be restored through mtDNA content mixing (Fig. 4d and Movie S7). Furthermore, considering that both IMMBright660 and SYBR™ Gold are uploaded by membrane potential [39], these results demonstrate that mitochondria lacking mtDNA but maintaining membrane potential can fuse with healthy mitochondria containing mtDNA to share mtDNA [40]. Therefore, fusion shares mtDNA through content mixing to preserve mitochondrial function.

We also observed the dynamic changes in cristae and mtDNA during fission (Fig. 4e and Movie S8). MtDNA was distributed in three parts of the mitochondrion. The initially narrow mitochondrion further shrunk at the beginning of fission. The results demonstrate that mitochondria were divided into three parts after fission, with each part containing a copy of mtDNA. Clearly, fission involved in cristae remodeling drives the distribution of mtDNA to subsequent daughter mitochondria.

By contrast, we also observed uneven dissemination of mtDNA during mitochondrial fission (Fig. 4f and Movie S9). Mitochondrial dysfunction leads to altered mitochondrial dynamics and selective removal of damaged mitochondria via mitophagy [41]. During fission, mtDNA on the top of the mitochondrion would be transferred to the bottom following cristae remodeling, resulting in the absence of mtDNA on the top of the mitochondrion. Loss of mtDNA may lead to the cessation of mitochondrial respiratory activity, which may be related to mitochondrial quality control and lipid metabolism.

MtDNA replication can be synchronized with fission to ensure the correct segregation of genetic material in most instances [36, 42]. We observed the entire process from mtDNA replication to mitochondrial division (Both IM and mtDNA imaging results were acquired by spinning disk confocal microscopy). As shown in a representative time-lapse series (Fig. 4g and Movie S10), this long mitochondrion has only one nucleoid (00:00 to 01:46). At 02:21, a nascent nucleoid appeared near the original nucleoid, then fission taked place between two nucleoids (02:46), and finally, the original mitochondria were divided into two daughter mitochondria, each with one nucleoid. Meanwhile, mtDNA was located at newly generated mitochondrial tips after fission. Our results directly demonstrate the spatial link between replicating mtDNA and fission sites. This mechanism allows each mitochondrion to obtain a copy of the genome following division and nucleoids are dispersed throughout the mitochondrial network.

We also observed that small mitochondrial branches detach from the mitochondrial network and fuse with nearby mitochondria (spinning disk confocal microscopy, Fig. 4h and 4i and Movie11). We assume that the small mitochondria function like a cargo, which can be separated from the mitochondrial network and transport mtDNA to other mitochondrial networks to drive the even distribution of mtDNA. We measured the area ratio in mitochondrial networks 1 and 2 as 0.21 and 0.12, respectively (white and blue dashed lines). The area ratio of mitochondrial network 2 fused with small mitochondria (area ratio 0.31, yellow dashed lines) reached 0.17, which was close to the average level, and the area ratio of mitochondrial network 1 after splitting was 0.18. Therefore, the fission and fusion of small mitochondria further promotes the even distribution of mtDNA, and tip distribution of mitochondria is more beneficial for the release of small mitochondria.

### Breakdown of cristae structure in apoptosis leads to mtDNA convergence

MtDNA-encoded proteins, such as components of ATP synthase, play a direct role in the formation and composition of cristae structure. ATP synthase is able to form dimers and oligomers, consisting of rows of dimers, which shape the mitochondrial IM and thus contribute to cristae formation [43]. Normal cristae structure ensures that mtDNA can be evenly distributed, since cristae can form partitioning structures to prevent the aggregation of mtDNA. On the contrary, once the cristae structure fails leading to mtDNA distribution disorder, mitochondria will face disaster. Remodeling of the cristae occurs during apoptosis to ensure the release of cytochrome c from the intracristal space into the cytoplasm, leading to downstream cascades of caspase activation and cell death [44]. During the prophase of apoptosis, the formation of higher-order oligomers of BAX/BAK creates lipid pores within the OM, resulting in IMM herniation and mtDNA efflux [45, 46].

After time-lapse STED imaging, the IM is remodeled during the process of apoptosis (Fig. 5a and Movie S12). We selected three mitochondrial cristae to measure the width change (white arrows 1, 2, and 3). The mitochondrial cristae gradually widened as the imaging frame number increased (Fig. 5b), implying that cytochrome c is released from the cristae lumen. This process was also accompanied by other cristae remodeling processes, including cristae merging and splitting and attachment to the inner boundary membrane in an arc, and this series of cristae remodeling processes may further promote apoptosis. MtDNA will gradually break free from the shackles of the mitochondrial cristae, and begin to converge, and the even distribution will be destroyed (Fig. 5a).

**Figure 5.**
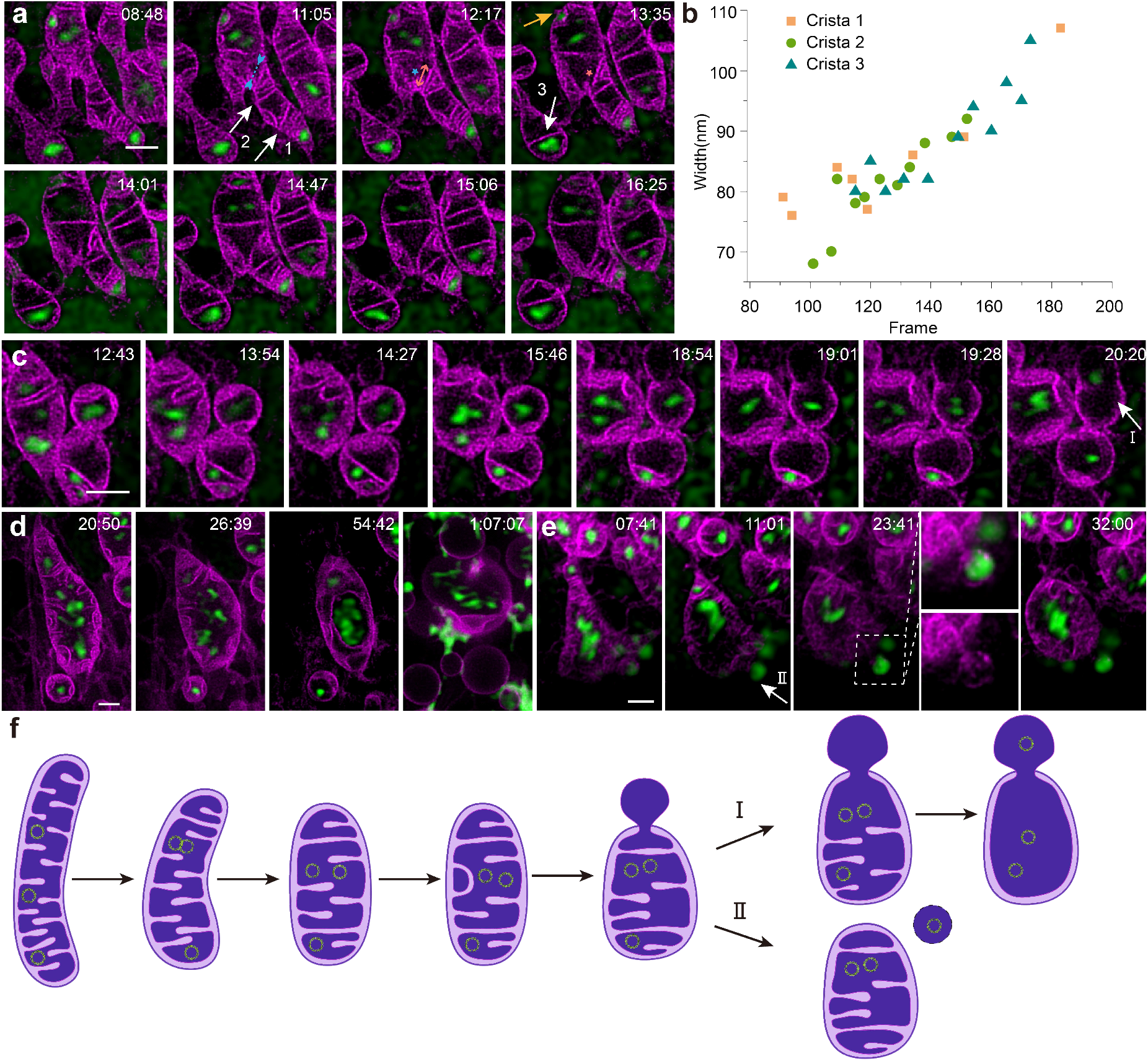
IM dynamics and mtDNA spatial changes during apoptosis. **a**, Cristae remodeling and mtDNA convergence during early apoptosis. The yellow arrow indicates that cristae are attached to the inner boundary membrane in an arc. Merging and splitting events of cristae are indicated by blue and orange asterisks, respectively, with bidirectional arrows facing inward (blue) and outward (orange) representing impending cristae merging and splitting events, respectively. **b,** Mitochondrial cristae gradually widened as the imaging frame number increased. The three curves represent the three cristae indicated by white arrows 1, 2, and 3 in Figure 5**a**, respectively. **c**, **d**, IMM herniation and mtDNA leakage along with cristae remodeling; white arrow 1 represents the first manner of mitochondrial herniation. **e**, The herniated IM forms single membrane-bound vesicles through budding off; white arrow 2 represents the second manner of mitochondrial herniation. **f**, Model of IM dynamics and mtDNA leakage during apoptosis. Scale bars in **a**, **c**, and **d** are 1 μm.

As reported [46], after BAK/BAX activation and loss of cytochrome c, large BAK/BAX pores appeared in the OM. These BAK/BAX macropores allowed the IM an outlet through which to herniate, carrying mtDNA. We observed the dynamic process of IMM herniation and mtDNA leakage along with cristae remodeling (Fig. 5c, Movie 13, Fig. 5d, Movie 14, Fig. 5e, Movie 15). The herniated IM formed a barbell-shaped structure, and mtDNA moved into the herniated IM, as shown by the white arrows in Fig. 5c. The herniated IM also formed single membrane-bound vesicles through budding off, as shown by the white arrows in Fig. 5e. During IM herniation, cristae within the mitochondria are pulled out, gradually disappearing (Fig. 5c). As the cristae within the mitochondria reduced, mtDNA was completely discharged from the tethering of mitochondrial cristae (Fig. 5c). Finally, the cristae structure and the distribution of mtDNA collapsed simultaneously.

### The spatial location of mitochondrial cristae and mtDNA during ferroptosis

Ferroptosis is an iron-dependent form of regulated necrosis in which intracellular reactive oxygen species (ROS) accumulation exceeds redox homeostasis that is maintained by glutathione (GSH) and phospholipid hydroperoxidase enzymes. The small-molecule compound erastin disrupts cellular redox homeostasis by inhibiting the cystine-glutamate antiporter, leading to depletion of cellular cysteine and glutathione [47, 48]. Morphologically, cells in ferroptosis exhibit mitochondrial ultrastructural changes such as decreased volume, disruption of the mitochondrial OM, and reduced mitochondrial cristae [49]. However, the manner by which the spatial conformations of cristae and mtDNA change under ferroptosis remains unknown.

We used erastin to induce ferroptosis in COS7 cells and performed imaging under STED. In comparison with the control, the mitochondria shrank into ellipsoids, and mitochondrial length and the number of cristae decreased (Fig. 6a,b). The distribution of mitochondrial length and the number of cristae in control and erastin-treated cells is shown in Fig. 6c. Statistical analysis shows that the majority of mitochondria in erastin-induced ferroptosis displayed shorter lengths and fewer cristae. We classified all mitochondria into three categories according to their length and number of cristae. Mitochondria with lengths shorter than 2.5 μm and less than 6 cristae were divided into Class I, those with lengths of 2.5–5 μm or 6–12 cristae were divided into Class II, and those with lengths longer than 5 μm or more than 12 cristae were divided into Class III. Representative mitochondria of these three classes in the control and erastin-treated groups are shown enlarged in Fig. 6a and 6b. We compared the three types of mitochondria and found that 55.7% of the mitochondria in the erastin-treated cells belong to Class I, while the majority of mitochondria in control cells belong to Class III (67.4%) (Fig. 6d). This indicates that erastin induces shrinking of mitochondrial morphology and a decrease in the number of cristae.

**Figure 6.**
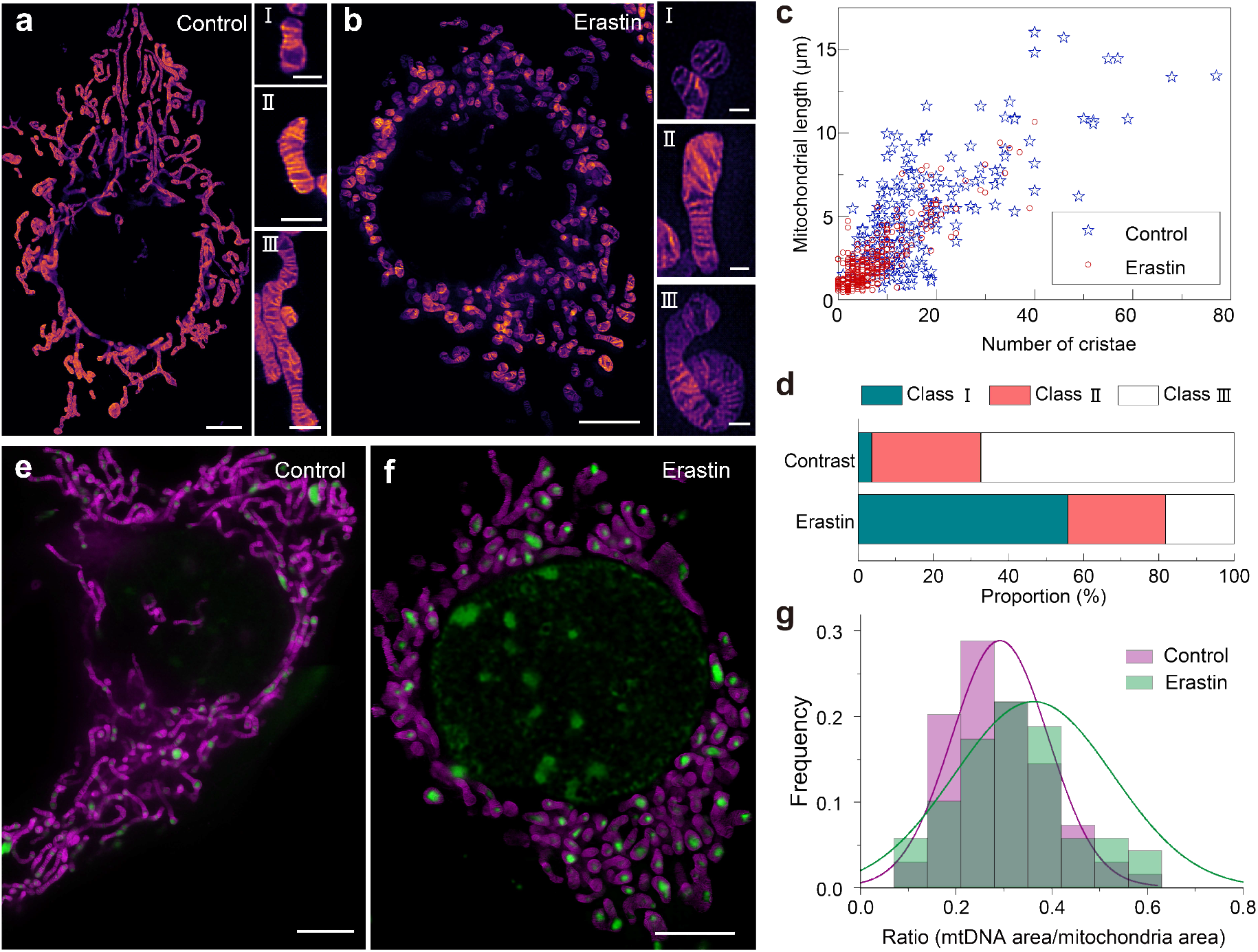
Spatial changes in mitochondrial cristae and mtDNA during ferroptosis by STED imaging. **a**, **b**, Representative mitochondrial cristae images of control and erastin-treated cells. Mitochondria were classified into three categories based on length and number of cristae, as shown on the right side of the panel. **c**, Plot of the mitochondrial length and number of cristae in control and erastin-treated cells. **d**, Proportion of the three types of mitochondria under different treatment groups. **e**, **f**, Dual-color live-cell imaging of IM and mtDNA in control and erastin-treated cells. **g**, Frequency distribution histogram of the ratio of mtDNA area to mitochondrial area. Scale bars in **a**, **b**, **e**, and **f** are 1 μm.

We performed dual-color imaging of mitochondrial cristae and mtDNA to observe the morphological changes during ferroptosis. We found that all mtDNA converges at the center of mitochondria, and the even distribution of mtDNA was disrupted (Fig. 6E and 6f). We calculated the ratio of the mtDNA area to the mitochondrial area in normal cells and those undergoing ferroptosis to plot the frequency distribution histogram (Fig. 6g). The area ratio in normal cells and those undergoing ferroptosis was 0.291 and 0.363, respectively. Statistical analysis shows no significant change in the area distribution of mtDNA between the control and erastin-treated groups, despite the significant morphological changes in the ferroptosis group (Fig. S7). We suppose that mitochondrial shrinkage may lead to an increase in the area ratio.

## Discussion

In this study, we developed a lipid membrane fluorogenic probe IMMBright660, which possesses the characteristics of high photostability, mitochondrial IM localization, and lipid membrane fluorogenicity. We used IMMBright660 and SYBR™ Gold to perform time-lapse dual-color STED/confocal super-resolution imaging of IM and mtDNA with excellent resolution. We found that mtDNA is regularly spaced within mitochondrial networks, preferring to be distributed at the tips and branch points. We observed the formation of the mitochondrial branch by fusion, and first visualized the cristae dynamics and the mtDNA changes in distribution when mitochondria extend the branch. According to statistical analysis, fusion always occurs near mtDNA. The void matrixes between cristae clusters where mtDNA locate are not dense, and we hypothesize that the pressure of cristae remodeling will be less when fusion occurs near mtDNA.

Mitochondria can adapt to various external changes, both as isolated entities and in solidarity with each other in vast sprawling networks [37]. Mitochondrial dynamics based on cristae remodeling may promote the even distribution of mtDNA. More specifically, mitochondrial cristae facilitate the rearrangement of mtDNA among different mitochondria during fusion and fission. Mitochondria may share mtDNA to preserve mitochondrial function, or unevenly disseminate mtDNA to daughter mitochondria for quality control. We also noticed that small mitochondria detach from the mitochondrial network, transport mtDNA to nearby mitochondrial networks, and further fuse with other mitochondria to drive the even distribution of mtDNA.

MtDNA plays an important role in maintaining cristae structure [43]. Conversely, maintenance of mtDNA distribution also requires normal cristae dynamics. The cristae structure fails in apoptosis and ferroptosis, leading to mtDNA distribution disorder. We found that cristae were remodeled to facilitate the release of cytochrome c, and the even distribution of mtDNA had been destroyed in apoptosis. We also captured the dynamic process of IM herniation and mtDNA leakage along with cristae remodeling in late apoptosis. We further observed mitochondria shrink into ellipsoids, mitochondrial length and the number of cristae decreased, and all mtDNA converged at the center of mitochondria during ferroptosis. We also propose the area ratio of mtDNA and mitochondrial, which can be used as a marker of mitochondrial function and status. Since metabolic activity is very low during both apoptosis and ferroptosis, we suggest that the decrease in mitochondrial cristae along with mitochondrial fragmentation may be an indicator of low metabolism.

In perspective, the direct association between mtDNA and the respiratory chain indicates that an even nucleoid distribution can achieve normal respiratory activity in the cell. On the contrary, an uneven nucleoid distribution signifies respiratory disaster or a change in the physiological state of the cell [10]. Form always follows function [6], and the dynamic layouts between the IM and mtDNA always respond to the mitochondrial function. Unfortunately, labeling of mtDNA limits the time-lapse STED imaging of mtDNA. In future studies, we intend to develop new mtDNA labeling methods that can be adapted to STED time-lapse imaging, and further explore the dynamic behavior of mitochondrial proteins involved in the interaction between the IM and mtDNA to elucidate the underlying mechanism [16, 50] with dual-color STED imaging.

## Materials and Methods

### Fluorescent probe design

Synthesis procedures for compounds and design strategies are shown in Supplementary Scheme S1. Chemical structure identification (^1^H, ^13^C NMR, and mass spectra) of compound **3** and **IMMBright660** are shown in Appendix.

### STED super-resolution for live cells

Cells were seeded on coverslips or glass-bottomed dishes and cultured to a suitable density (24 h) at 37 °C in a 5% CO2 atmosphere with 95% humidity. Cells were incubated with IMMBright660 for 10 mins under the same conditions, after which images were obtained using a confocal laser-scanning microscope. STED imaging was performed using the Abberior Facility Line (Abberior Instruments GmbH, Germany) with a 637-nm laser for excitation and a 775-nm pulsed laser for STED depletion. A 60× oil-immersion objective (N.A. 1.42, Olympus, Japan) was employed in imaging experiments. All STED results presented in the paper have been processed by deconvolution software (Huygens, SVI, Netherlands). The deconvolution parameter is set to improve the resolution while maintaining the real structure.

### Spinning-disk confocal microscopy

Spinning disk confocal data were acquired under a yokogawa spinning disk equipped with a Live SR super resolution module (Gataca systems, France), enabling a 2x resolution improvement at very high imaging speeds. A 100× oil-immersion objective (N.A. 1.4, Niokn, Japan) was used in live cell imaging.

### Segmentation of mitochondrial cristae and statistical analyses

The areas of mitochondrial cristae and mtDNA were calculated using ImageJ plugins. We applied the ImageJ plugin Trainable Weka Segmentation to train the cristae classifier and obtain probability images [51]. The threshold was adjusted until cristae were accurately distinguished from the background, and binary images were then obtained. For binary images, mitochondrial and mtDNA areas were calculated using the analysis function in ImageJ.

## Contributions

P.X. and B.G. initiated the project. W.R. and B.G. conceived the experiments. B.G., X.G., S.L., and W.R. designed and measured the probe properties. C.S. and P.X. provided imaging conditions. W.R., X.G., and C.S. performed imaging experiments. W.R. and X.G. analyzed the imaging data. M.L. provided cell culture guidance and conducted effective discussions. P.X., B.G., and C.S. supervised the project. X.G. and W.R. prepared the figures. W.R., X.G., C.S., P.X., and B.G. wrote the manuscript with contributions from all authors.

## Acknowledgments

This work was supported by the National Key R&D Program of China (2022YFC3401100), and National Natural Science Foundation of China (22177024, 62025501, 31971376, 92150301). We thank the National Center for Protein Sciences at Peking University in Beijing, China, for assistance with STED super-resolution imaging. We thank abberior China for their guidance in measuring saturation intensity and secondary excitation effects, and Optofem Technology Limited for providing Facility Line STED and Huygens software. We thank Dr. Peiyuan Chai from Prof. Junlin Teng’s Lab for providing HeLa cells and Dr. Huiwen Hao from Prof. Yujie Sun’s Lab for providing COS7 cells. We would like to thank Professor Congying Wu and Professor Bo Li for their helpful discussions and suggestions.

