## Supporting Information (SI) for "Visualization of mitochondrial cristae and mtDNA evolvement and interactions with super-resolution microscopy"

**This PDF file includes:**

Supplementary methods
Supplementary chemical synthesis
Supplementary Table S1
Figures S1 to S7
NMR spectra of newly synthesized compounds and Mass spectra analysis

**Other supporting materials for this manuscript include the following:**

Movies S1 to S15

**Supplementary methods**

**UV/Vis absorption and fluorescence spectroscopy measurements.** Si-rhodamine probe was dissolved in DMSO to prepare a stock solution (10 mM) and used at a working concentration of 10 μM. UV/Vis absorption spectra of IMMBright660 were measured using an Analytikjena Specord 210 plus spectrophotometer (Analytikjena, German), and fluorescence emission spectrum was measured with Hitachi F-7000 fluorescence spectrophotometer (Hitachi, Japan). All measurements were performed in a 3-mL cuorvette with 2 mL solution. The absorption and fluorescence spectra of probe IMMBright660 (10 μM) were measured in the sodium chloride solution (150 μM) and Artificial lipid membrane solution (2 mM). Dimyristyl phosphatidyl choline (DMPC) (32.3 mg) and dimyristyl phosphatidyl glycerophosphate (DMPG) (8.21 mg) were dissloved in 20 mL mixture solution (MeOH/DCM = 1/4, v/v). After evaporating solvent and drying for 2 h under reduced pressure, the solid was dissolved in 28.9 mL sodium chloride solution (100 mM), further passed argon for 10 min to remove the dissolved oxygen in the solution, ultrasonic 5 min, passed aqueous polycarbonate membrane (200 nm) for 21 times.

**Fluorescence quantum yield measurements**. The fluorescent quantum yields for Probe Si-rhodamines were measured in DMSO using Alexa Fluor 647 (Φ = 0.27, in PBS) as the standard substance at an excitation wavelength of 646 nm, and the quantum yields were calculated using the following equation: $\Phi_s = \Phi_r (A_rF_s/A_sF_r)(n_s^2/n_r^2)$, where s and r denote sample and reference, respectively, A is the absorbance, F is the relative integrated fluorescence intensity, and n is the refractive index of the solvent.

**Photon-stability measurements.** The photon-stability of probe IMMBright660 (10 μM) and Alexa Fluor 647 in different solution including $H_2O$, NaCl solution (10 mM) and artificial lipid membrane solution (2 mM) were irradiated with a 1 W 660 nm LED laser (1 W on sample). Absorption spectra were measured after irradiation, and relative absorbance at the maximum wavelength was plotted as a function of irradiation time with a laser.

**Cell culture.** COS7 cells were cultured at a suitable density (moderate) in DMEM supplemented with penicillin (100 units/mL), 10% (v/v) FBS (WelGene), and streptomycin (100 mg/mL) at 37°C in a 5% $CO_2$ atmosphere with 95% humidity. Cells were seeded on confocal dishes 24 h prior to the experiments.

**Experimental Procedures**

**Materials and characterization.** All chemicals used for synthesis were purchased from commercial suppliers and applied directly in the experiment without further purification. Solvents were either employed as purchased or dried according to procedures described in the literature. The progress of the reaction was monitored by TLC on pre-coated silica plates (GF-254, 250 μm in thickness), and spots were visualized by UV light. Qingdao ocean silica gel (100-200 mesh) was used for general column chromatography purification. $^{1}H$ NMR and $^{13}C$ NMR spectra were recorded on Q. One AS 400 or Bruker 600 spectrometer with $CDCl_3$, DMSO-$d_6$ or $CD_3OD$ as solvent. Chemical shifts are reported in parts per million relative to internal standard tetramethylsilane ($Si(CH_3)_4$ = 0.00 ppm). High-resolution mass spectra (HRMS) were obtained on a XEVO-G2QTOF (ESI) (Waters, USA) or TSQ Quantum Ultra (ESI) (Thermos, Germany). The UV-visible spectra were recorded on a Specord 210 plus spectrophotometer (Analytikjena, Germany). Fluorescence study was carried out using F-7000 fluorescence spectrophotometer (HITACHI, Japan) at 25°C. Dynamic light scattering (DLS) was determined with Winner 802 (Jnwinner, China).

**Synthesis**

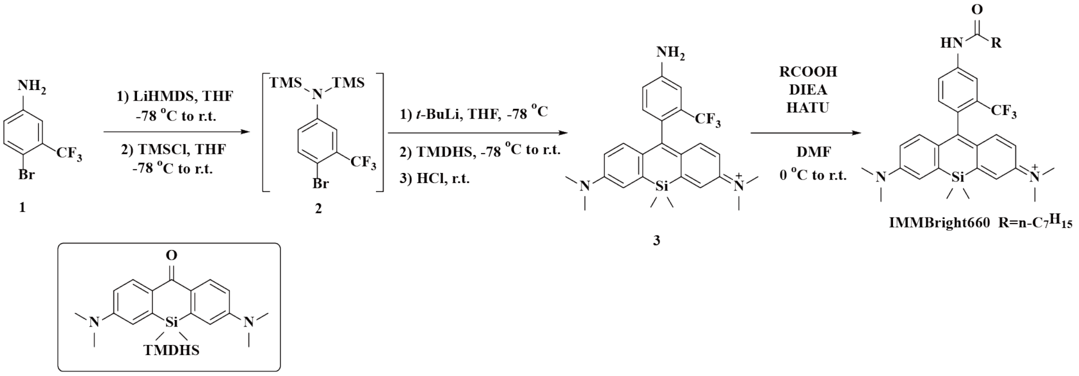


**Scheme. S1.** Paste figure above the legend Synthetic scheme for Si-rhodamines

**Compound 2.** In a nitrogen-flushed flask fitted with a double port reaction bottle, compound **1** (360 mg, 1.5 mmol) was dissolved in anhydrous THF (10 mL) and the solution was cooled to -78 °C. 1.3 M Lithium bis(trimethylsilyl)amide (LiHMDS) 2.5 mL, 3.3 mmol) was slowly added dropwise via a syringe to the above solution in an $N_2$ atmosphere. After that, the reaction solution was stirred for 20 minutes at -78 °C, then warmed to room temperature, and further stirred for 5 min. After further cooling to -78°C, dimethyldichlorosilane (TMSCl) (359 mg, 3.3 mmol) dissolved in anhydrous THF was slowly added into the system, then the solution was warmed to room temperature and stirred for 16 h. The solvent was evaporated at reduced pressure to obtain intermediate **2** without separation, which was used directly for the next step.

**Compound 3.** In a nitrogen-flushed flask fitted with a double port reaction bottle, intermediate **2** was dissolved in anhydrous THF (10 mL) and the solution was cooled to -78 °C. 1.3 M *t*-BuLi (1.1 mL, 1.5 mmol) was slowly added dropwise via a syringe to the above solution in an $N_2$ atmosphere. After that, the reaction solution was stirred for 30 minutes at -78°C. TMDHS (50 mg, 0.15 mmol) dissolved in anhydrous THF was slowly added into the system, then the solution was warmed to room temperature and stirred for 2 h, quenched with 2 M HCl. The aqueous solution was extracted with dichloromethane, and the combined organic phase was washed with water and brine, dried over $Na_2SO_4$, filtered and evaporated. The resultant residue was quickly purified by silica gel chromatography (MeOH/DCM = 1/50, v/v), yielded 74%, 53 mg of pure product as blue solid. $^1$H NMR (400 MHz, $CDCl_3$) $\delta$ 7.14 (d, $J$ = 10.2 Hz, 2H), 7.09 (d, $J$ = 8.7 Hz, 4H), 6.84 (d, $J$ = 8.2 Hz, 1H), 6.56 (dd, $J$ = 9.6, 2.3 Hz, 2H), 3.34 (s, 12H), 0.59 (s, 3H), 0.46 (s, 3H). $^{13}$C NMR (101 MHz, $CDCl_3$) $\delta$ 168.94, 153.89, 148.80, 148.10, 142.40, 131.47, 128.81, 125.14, 123.77, 122.41, 120.26, 117.43, 113.49, 111.83, 77.48, 77.16, 76.84, 40.99, 29.61, -0.27, -1.93. MALDI-TOF MS m/z Calculated 468.2077 for $C_{26}H_{29}F_3N_3Si^+$, found 468.1682 $[M]^+$.

**IMMBright660**. In a flame-dried flask flushed with nitrogen, n-octylic acid (17.3 mg, 0.12 mmol) was dissolved in anhydrous DMF (10 mL). HATU (55 mg, 0.144 mmol) was added at ice bath. After stirred at room temperature for 0.5 h, compound **3** (28 mg, 0.06 mmol) and DIPEA (31 mg, 0.24 mmol) were added one by one at ice bath. After further stirred for 3 h, the reaction solution was extracted with dichloromethane. The organic layer was dried over $Na_2SO_4$ and evaporated. The resultant residue was quickly purified by silica gel chromatography (MeOH/DCM = 1/100, v/v), yielded 84%, 30 mg of pure product as blue solid. $^1$H NMR (400 MHz, DMSO-$d_6$) $\delta$ 10.47 (s, 1H), 8.30 (s, 1H), 7.98 (d, $J$ = 8.3 Hz, 1H), 7.42 (s, 2H), 7.35 (d, $J$ = 8.2 Hz, 1H), 6.84 (q, $J$ = 9.7 Hz, 4H), 3.31 (s, 12H), 2.40 (t, $J$ = 7.0 Hz, 2H), 1.64 (s, 2H), 1.30 (d, $J$ = 14.0 Hz, 8H), 0.90 - 0.83 (m, 4H), 0.64 (s, 3H), 0.51 (s, 3H). $^{13}$C NMR (101 MHz, $CDCl_3$) $\delta$ 173.88, 167.56, 154.45, 148.78, 142.68, 140.42, 131.65, 131.22, 130.35, 128.77, 122.99, 120.90, 118.22, 114.12, 77.80, 77.48, 77.16, 41.29, 37.72, 32.37, 32.18, 30.15, 27.66, 25.95, 23.10, 14.58, 0.04, -1.73. MALDI-TOF MS m/z Calculated 594.3122 for $C_{34}H_{43}F_3N_3OSi^+$, found 594.3130 $[M]^+$.

**Supplementary Fig. S1 to S7**

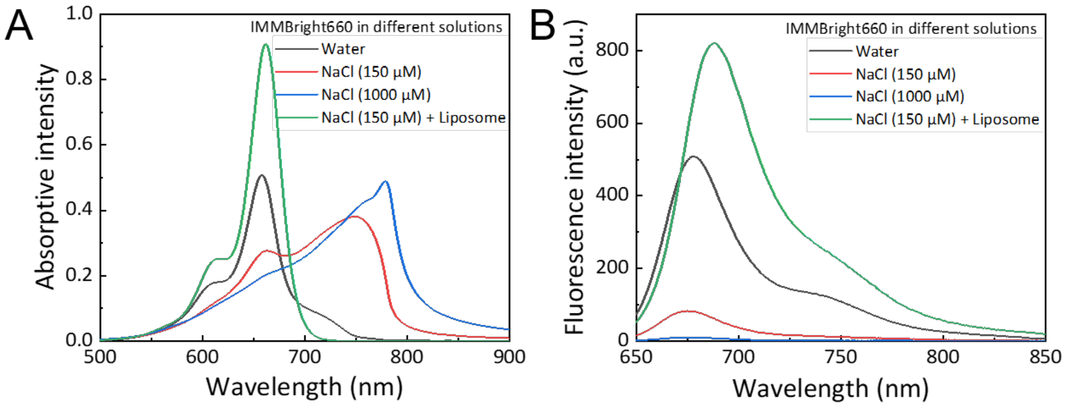


**Fig. S1.** Absorption (*A*) and fluorescence (*B*) spectra of IMMBright660 solution in NaCl solution in the presence or absence of liposome.

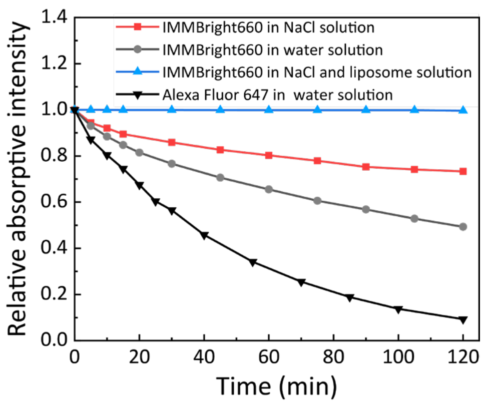


**Fig. S2.** The particle size distribution of IMMBright660 in different concentrations of NaCl solution.

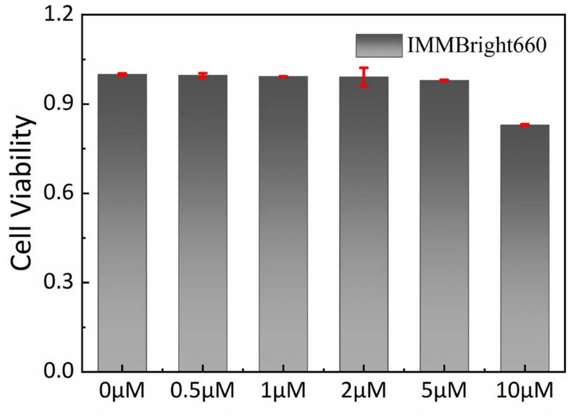


**Fig. S3.** The cytotoxicity of probe IMMBright660 using a standard MTT assay.

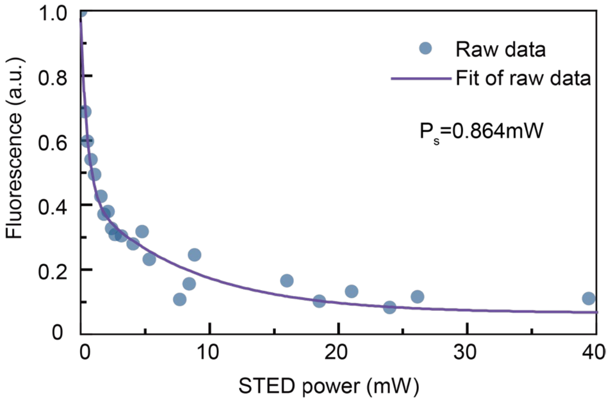


**Fig. S4.** The detected fluorescence signal of the IMMBright660 solution dye pool on a coverslip as a function of the depletion beam intensity; the excitation beam was 660 nm.

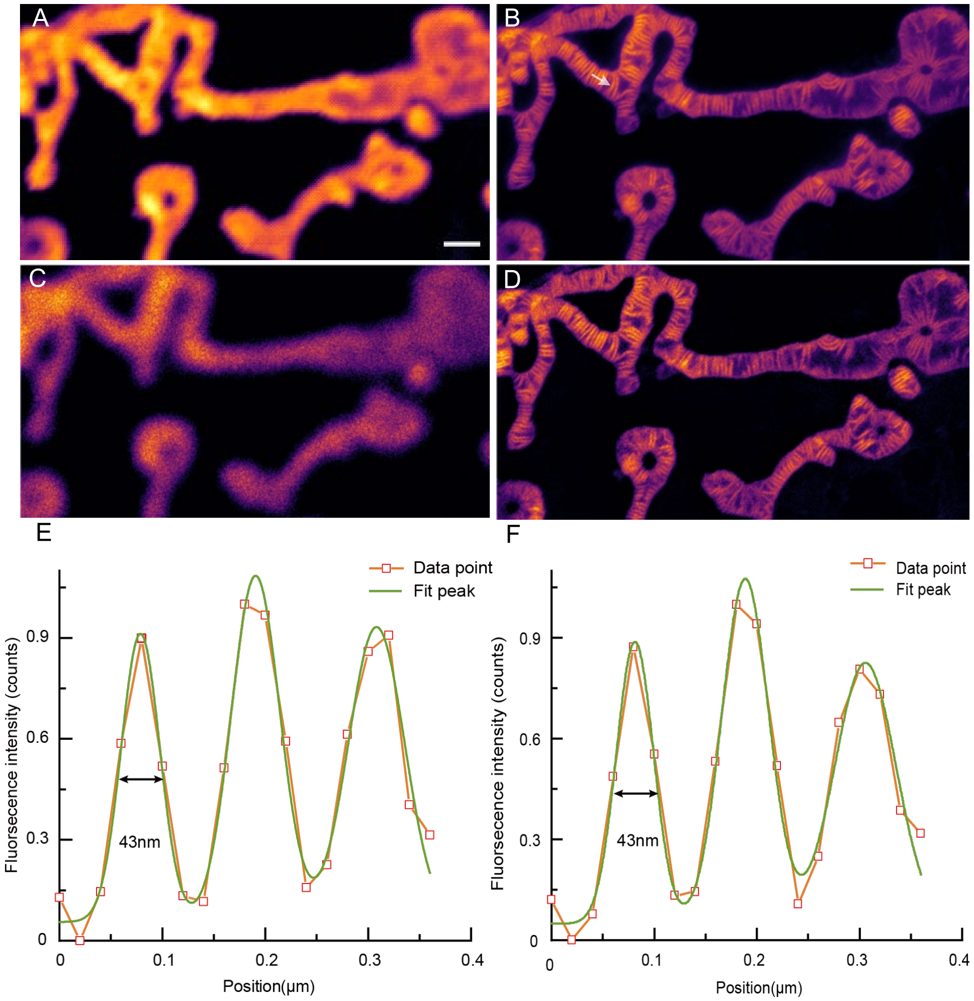


**Fig. S5.** Confocal (*A*) and STED (*B*) images of living COS7 cell mitochondria labeled with IMMBright660. (*C*) Mitochondria of COS7 live-cell were excited by depletion beam. (*D*) Signal of fig (*B*) subtracted signal of fig (*C*). (*E*) The signal intensity profile crossed the cristae (indicated with arrow) in fig (*B*). (*F*) The signal intensity profile crossed the cristae (indicated with arrow) in fig (*D*).

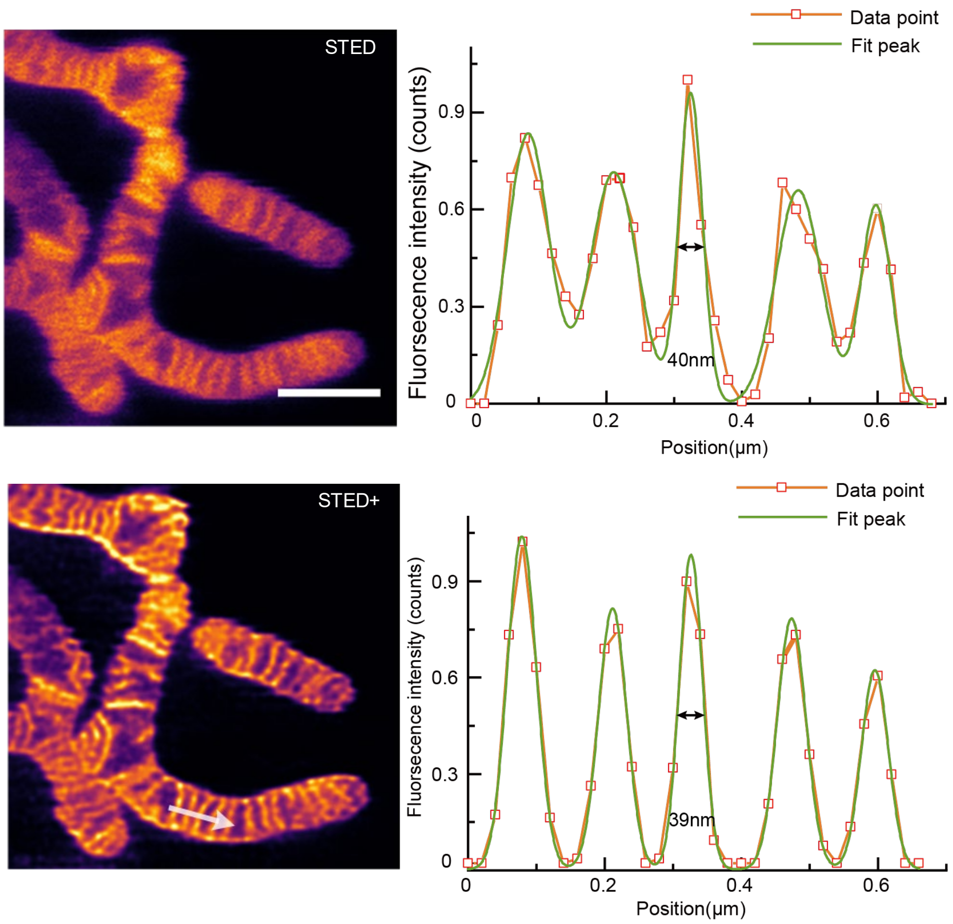


**Fig. S6.** STED and STED+ images(left) of mitochondria in live COS7 cells. The signal intensity profile (right) crossed the cristae (indicated with arrow).

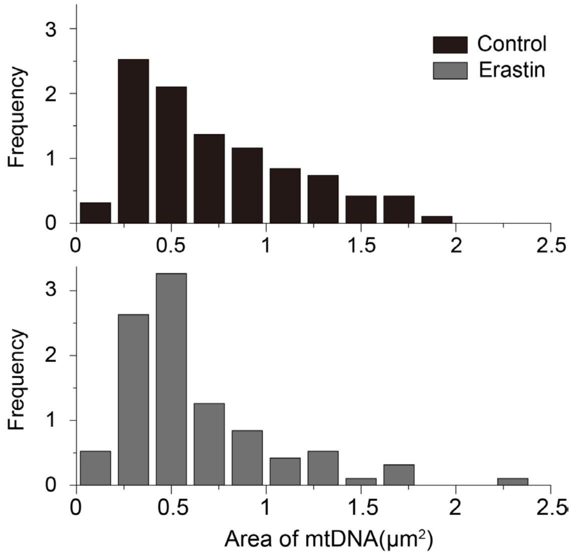


**Fig. S7.** Frequency distribution histogram of mtDNA area between the control and erastin-treated groups

**Captions for movies S1 to S15**

**Movie S1.** Time-lapse STED imaging of mitochondria labeled with IMMBright660.

**Movie S2.** Time-lapse STED imaging of spatial changes in mtDNA and cristae at the tip.

**Movie S3.** Branch point formation by mitochondrial fusion.

**Movie S4.** Branch point formation by emersion of a new branch.

**Movie S5.** The mtDNA on the mitochondrial branch moves to the branch point.

**Movie S6.** Time-lapse STED imaging of the mitochondrial fusion.

**Movie S7.** Fusion of a mitochondrion containing mtDNA and another mitochondrion without mtDNA.

**Movie S8.** Time-lapse STED imaging of the mitochondrial fission.

**Movie S9.** Mitochondria after fission may not contain mtDNA

**Movie S10.** MtDNA replication initiates mitochondrial division

**Movie S11.** Small mitochondrial branches detach from the mitochondrial network and fuse with nearby mitochondria.

**Movie S12.** Cristae remodeling and mtDNA convergence during early apoptosis.

**Movie S13.** IMM herniation and mtDNA leakage along with cristae remodeling. The progress represents the first manner of mitochondrial herniation

**Movie S14.** IMM herniation and mtDNA leakage.

**Movie S15**. The herniated IM forms single membrane-bound vesicles through budding off. The progress represents the second manner of mitochondrial herniation

**NMR spectra of newly synthesized compounds and Mass Spectra analysis**

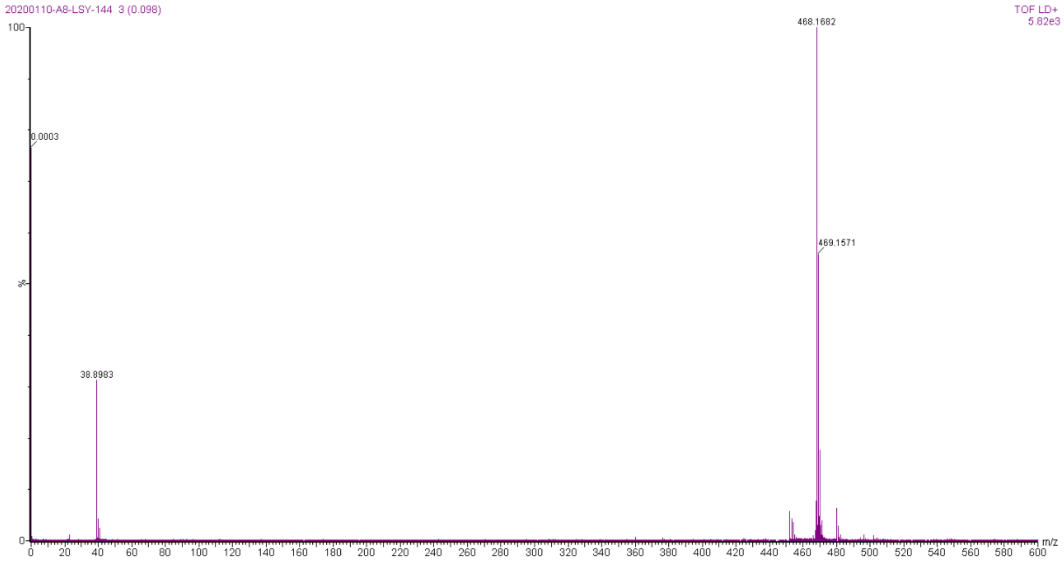


**MS-1.** MALDI-TOF-MS spectra of compound 3

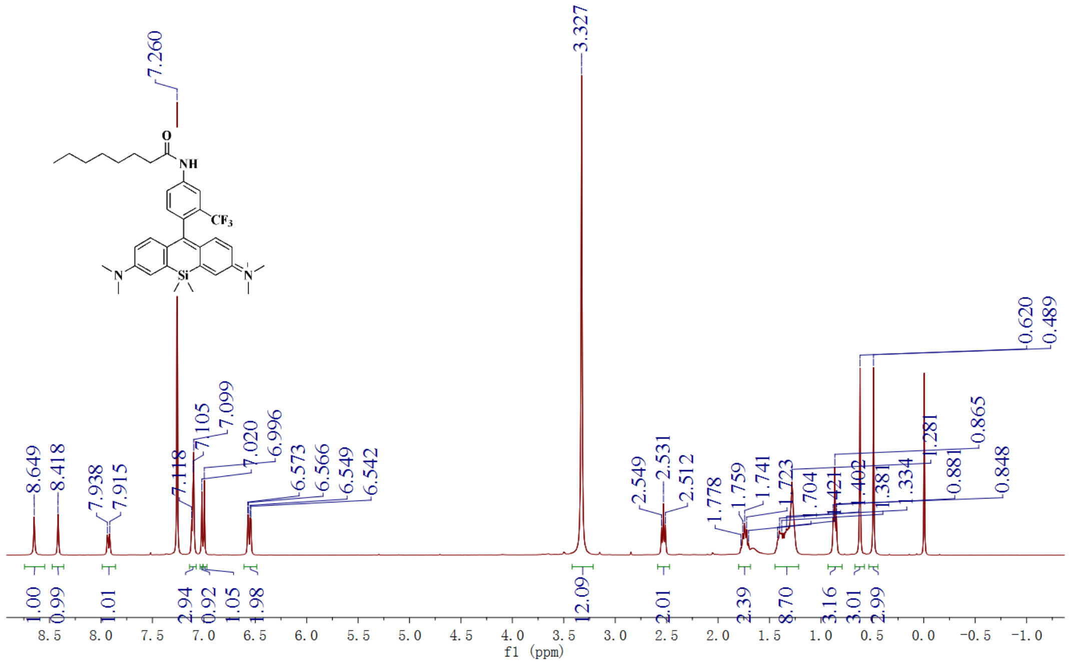


**NMR-1.** $^1$H NMR (400 MHz) spectra of IMMBright660 in $CDCl_3$.

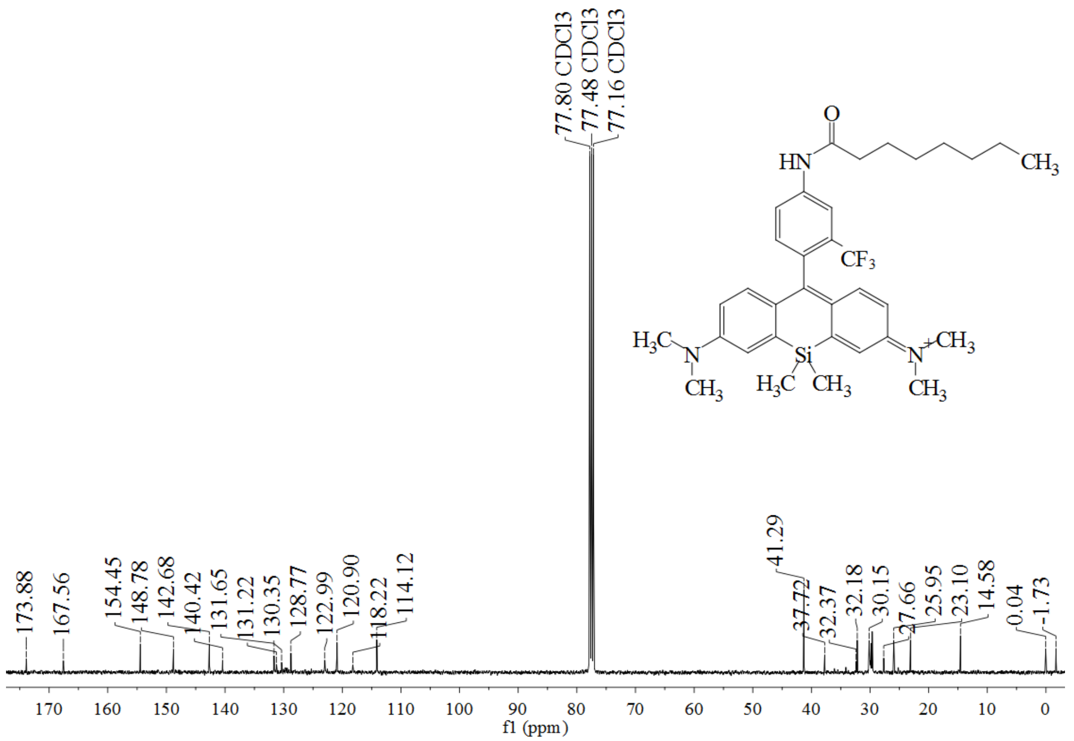


**NMR-2.** $^{13}$C NMR (101 MHz) spectra of IMMBright660 in $CDCl_3$.

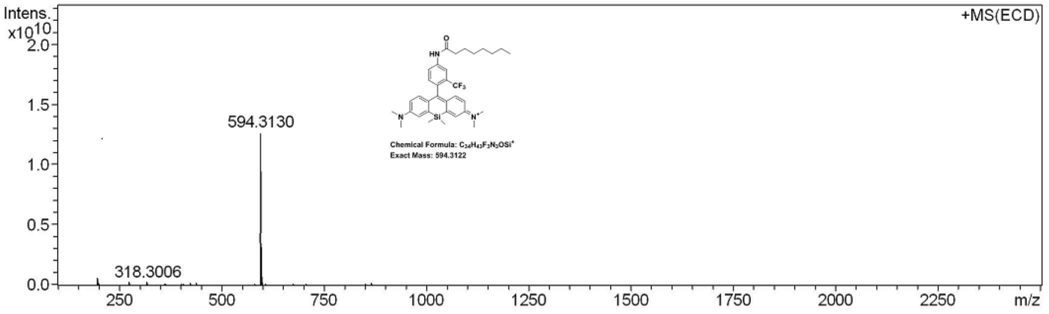


| Meas. m/z | # | Formula | Score | m/z | err [mDa] | err [ppm] | mSigma | rdb | e⁻ Conf | N-Rule |
|---|---|---|---|---|---|---|---|---|---|---|
| 594.3130 | 1 | C 34 H 43 F 3 N 3 O Si | 100.00 | 594.3122 | -0.8 | -1.4 | 27.3 | 14.5 | even | ok |

**MS-1.** High Resolution Mass Spectra of IMMBright660.
